# Non-toxigenic *Vibrio cholerae* challenge strains for evaluating vaccine efficacy and inferring mechanisms of protection

**DOI:** 10.1101/2021.12.17.473008

**Authors:** Bolutife Fakoya, Karthik Hullahalli, Daniel H. F. Rubin, Deborah R. Leitner, Roma Chilengi, David A. Sack, Matthew K. Waldor

## Abstract

Human challenge studies are instrumental for testing cholera vaccines, but these studies use outdated strains and require inpatient facilities. Here, we created next-generation isogenic Ogawa and Inaba *V. cholerae* challenge strains (ZChol strains) derived from a contemporary Zambian clinical isolate representative of current dominant pandemic *V. cholerae*. To minimize the risk of severe diarrhea these strains were rendered non-toxigenic, since antibody responses which limit *V. cholerae* colonization are the primary mechanism of immune protection. These strains did not cause diarrhea in infant mice and proved to accurately gauge reduction in intestinal colonization mediated by effective vaccination. They are also valuable as targets for measuring vibriocidal antibody responses. Using barcoded ZChol strains, we discovered that vaccination tightens the infection bottleneck without restricting pathogen expansion *in vivo*. ZChol strains have the potential to enhance the safety, relevance, and scope of future cholera vaccine challenge studies and be valuable reagents for studies of immunity to cholera.

## Introduction

Diarrheal diseases remain one of the leading causes of infectious disease death globally and cholera, caused by the human bacterial pathogen *Vibrio cholerae*, accounts for approximately 100,000 deaths each year^1,2^. Cholera is endemic in over 50 countries, and often spreads explosively during epidemics^1^ by the fecal-oral route and/or through ingestion of contaminated food or water. *V. cholerae* replicate in the human small intestine (SI) and secrete cholera toxin (CT), an AB5 type exotoxin that causes profuse secretory diarrhea, the clinical hallmark of cholera^3^. Rehydration therapy is the mainstay of cholera treatment with antibiotics being used in severe clinical cases. Several vaccine formulations have been developed to prevent cholera and three killed whole cell oral vaccine preparations are WHO prequalified, two of which, Euvichol and Shanchol, are distributed through a global stockpile^4,5^. It is thought that eradicating cholera globally will require multi-sectoral public health approaches, including vaccines, enhanced surveillance, as well as global water, sanitation, and hygiene initiatives^6^.

Though there are over 200 serogroups of *V. cholerae* only serogroup O1 has given rise to cholera pandemics^3^. Serogroup O1 *V. cholerae* are further classified into Ogawa and Inaba serotypes that differ in the methylation of the terminal perosamine of the lipopolysaccharide (LPS) O-antigen; Ogawa strains are methylated and Inaba strains are unmethylated^7^. Biotype is another key classifier of pandemic *V. cholerae*; the first six pandemics were caused by the now extinct classical biotype^8^, whereas the ongoing 7th pandemic is caused by the El Tor biotype^3^. Molecular epidemiology has identified 3 distinct clades (waves) of pandemic *V. cholerae* within the 7th pandemic that have emerged and disseminated from south Asia to other continents. Wave 3 7th pandemic El Tor (7PET) strains are responsible for the dominant circulating *V. cholerae* globally and have caused devastating outbreaks in Haiti, Yemen and in South Asia^9–11^. Currently, Sub-Saharan Africa carries the dominant burden of cholera cases reported to the WHO^12^.

Both animal and human studies suggest that immune responses targeting *V. cholerae* LPS are important for future protection from cholera, but responses to other antigens, including CT can also contribute to protection. Studies of cholera patients as well as healthy volunteers challenged with wild type *V. cholerae* revealed that infected persons develop serum and intestinal antibody responses to LPS and CT and were protected from future episodes ^13–15^. These observations motivated development of vaccines designed to stimulate immune responses similar to those stimulated by the disease^16^ and led to the development of killed whole cell oral vaccines ^3^ as well as a live attenuated oral vaccine^17^. Serologic correlates of protection are often useful to predict whether disease or vaccination induces protection^18,19^. For cholera, the vibriocidal antibody titer (VAT) response following vaccination does suggest protection, but is not a true correlate because VAT titers fall within a few months after vaccination, but protection lasts for several years^13,15,20^. Thus, proof of vaccine efficacy has depended on testing of vaccines using human volunteers or in controlled field trials.

Following a controlled human infection model (CHIM) showing that a killed whole cell oral cholera vaccine provides protection^21^, large, double-blind, placebo-controlled field trials demonstrated their efficacy in India^20^ and Bangladesh^22^ and subsequent case control studies confirmed its effectiveness in Africa^23^. A live attenuated vaccine was licensed for travelers based on results from a CHIM study that demonstrated protection for at least 3 months^24^, showcasing how CHIM studies have played a crucial role in cholera vaccine development. In these studies, the rate and severity of diarrheal illness in vaccinated and naïve volunteers who orally consume virulent *V. cholerae* is compared^24,25^. Typically, most of the unvaccinated, naïve volunteers develop diarrhea and/or vomiting for one to three days and require rehydration. Because of the risk of illness, cholera CHIMs are only carried out in specialized inpatient facilities where healthy volunteers are cared for by experienced physicians. CHIM studies for cholera were initially carried out by Cash et. al. to determine *V. cholerae* infectious doses and evaluate vaccine efficacy^14^. CHIM studies have also revealed that antibacterial immune responses lead to reductions in fecal excretion of challenge strains which generally correlate with vaccine efficacy^25^.

Currently, the *V. cholerae* strain used in most CHIMs is N16961, a 1971 7PET wave 1 Inaba isolate^24–27^. Although N16961 has been widely used for CHIM studies there are several compelling reasons why a new challenge strain would be valuable for cholera CHIM studies. First, N16961 is a wave 1 *V. cholerae* strain that differs considerably from currently circulating wave 3 *V. cholerae*, including in virulence associated loci such as *tcpA* and *ctxB^9^*. Second, N16961 strain exhibits hypervirulence in preclinical animal models of infection^28^. Third, the supply of Good Manufacturing Practice (GMP) manufactured N16961 that can be used in CHIM studies has become very limited. Fourth, since many of the volunteers develop severe diarrhea and vomiting, a CHIM strain that does not cause severe illness and without the need for a specialized center would be preferable if it provides evidence for vaccine protection. Given the potential risks to volunteers and biosafety concerns associated with toxigenic *V. cholerae*,cholera CHIM studies have not been carried out in cholera endemic regions^29,30^.

CHIMs have played a vital role in guiding cholera vaccine development, and since N16961 has significant limitations, we engineered *V. cholerae* strains, as potential next generation *V. cholerae* strains for volunteer challenge studies. These ZChol strains are derived from a 2016 7PET wave 3 clinical isolate from Zambia. To reduce the risk for volunteers and to potentially enable their conduct in cholera endemic countries, we rendered the strains non-toxigenic through deletion of *ctxAB* (CT). Isogenic Ogawa and Inaba versions of ZChol were also generated and these strains colonize the intestine of infant mice to similar levels as the wild type toxigenic parent strain (ZTox). We show that these strains robustly report on vaccine mediated protection both by colony-forming unit (CFU) reduction and as VAT targets. Finally, using ZChol strains barcoded with thousands of genomic sequence tags, we demonstrate the utility of barcoded and non-toxigenic challenge strains to decipher mechanisms of vaccine protection. Together these results indicate that the ZChol strains will facilitate an expanded range of safer *V. cholerae* CHIMs as well as laboratory studies of immunity to *V. cholerae*.

## Results

### ZTox is a wave 3 seventh pandemic El Tor *V. cholerae* strain

The *V. cholerae* strain that is most frequently used in challenge studies for testing vaccine efficacy, N16961, was isolated five decades ago and is no longer representative of the dominant globally circulating *V. cholerae*. We chose a 2016 *V. cholerae* O1 Ogawa clinical isolate from Zambia for development as a new challenge strain, reasoning that this isolate would be representative of contemporary pandemic *V. cholerae*. A combination of long and short read next generation sequencing (nanopore and Illumina, respectively) was used to compile this strain’s genome and determine where it falls within a representative phylogeny of toxigenic *V. cholerae* that was created using ~1200 clinical *V. cholerae* isolates^31^, including a substantial collection of contemporary African isolates. Comparison of ZTox with known toxigenic *V. cholerae* clinical isolates revealed that it clusters with 7PET wave 3 strains (Fig. 1), the dominant cause of cholera worldwide today. Genomic analyses of clinical *V. cholerae* isolates from Africa have revealed multiple re-introduction events of *V. cholerae* onto the continent via long-range transmission from South Asia^32^. The most recent of these events, which gave rise to the T13 sublineage, originates from the same clade that was introduced from Nepal to Haiti and gave rise to the Haitian cholera epidemic that began in late 2010 ^33^. ZTox is found within this T13 sublineage and clusters with recent outbreak strains from Southern and East Africa^34^. ZTox is also closely related to the *V. cholerae* strains that originated the devastating outbreak in Yemen throughout the late 2010^33^. These analyses suggest that this isolate is typical of present-day clinically relevant wave 3 El Tor *V. cholerae*.

**Figure 1.**
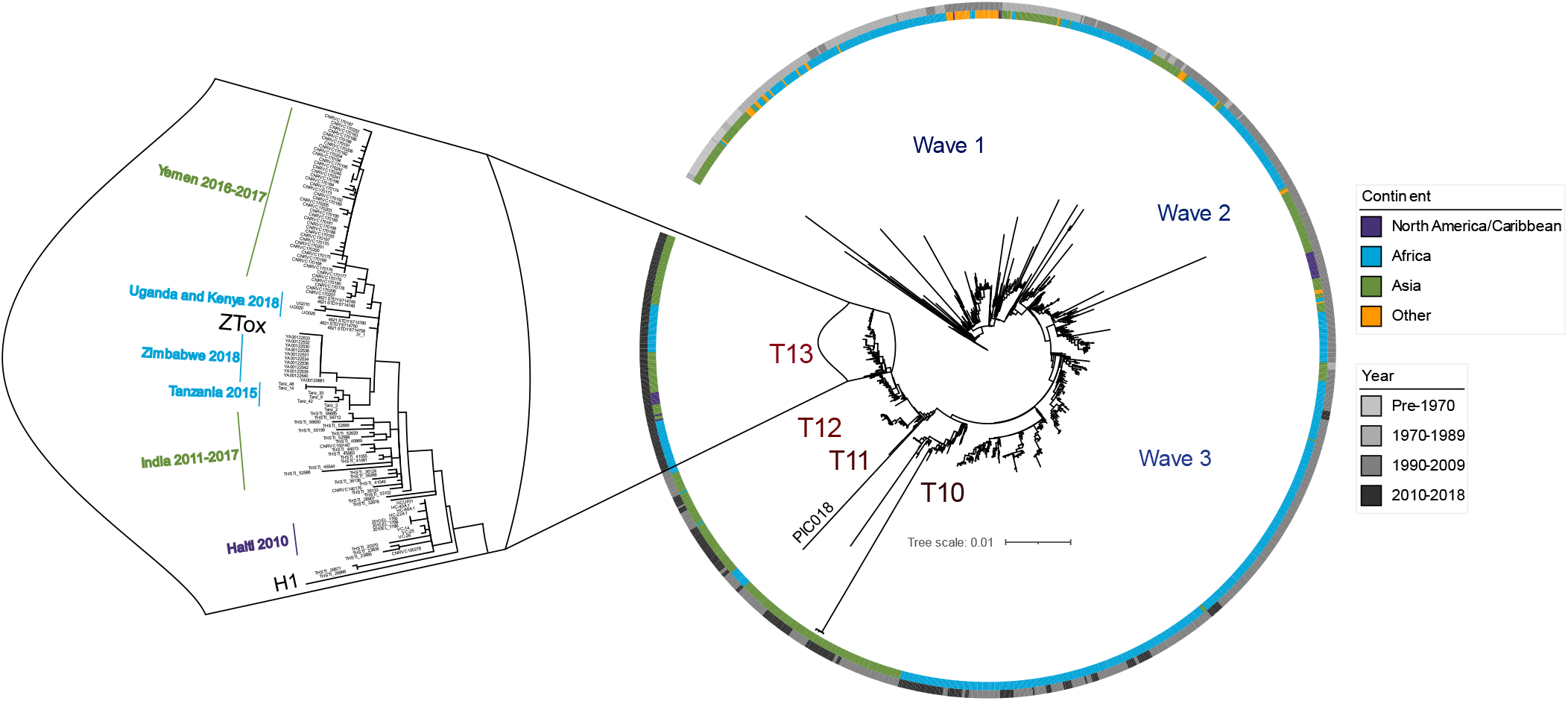
Position of ZTox in the phylogeny of pandemic *V. cholerae*. A maximum-likelihood phylogenetic tree from ~1200 toxigenic clinical isolates of *V. cholerae* with overrepresentation of contemporary clinical isolates from outbreaks in Sub-Saharan Africa (light blue). The pre-seventh pandemic strain A6^31^ was used as an outgroup. Scale bar represents the mean number of nucleotide substitutions per site. ZTox clusters with 7PET wave three strains, particularly with isolates from nearby African countries (sublineage T13) and South Asian *V. cholerae* endemic regions that seed long-range transmission events. The PIC018 Inaba strain used for vibriocidal assays is also depicted.

Analyses of important mobile elements within the ZTox genome also confirmed it is representative of contemporary wave 3 El Tor *V. cholerae*. The CTXΦ prophage in ZTox contains a single CTXΦ prophage and an adjacent RS1 satellite prophage^35^. Both elements encode the El Tor CTX phage repressor (rstR^ET^) (Fig. S1A). Like other wave 3 T13 strains, the ZTox CTX prophage harbors the *ctxB7* allele that has been associated with elevated secretion of cholera toxin (Fig. S1A). The SXT integrative conjugative element (ICE) commonly confers resistance to several antibiotics including sulfamethoxazole and low levels of chloramphenicol and streptomycin, and has been linked to evolution of both wave 2 and 3 7PET *V. cholerae^11^*. However, like many other recent African T13 strains, ZTox is sensitive to these antibiotics. Notably, the ZTox-SXT lacks *sul2*, and *floR* and *aph3*, and *aph6* that confer resistance to these agents (Fig. S1B) and is nearly identical to the SXT/R391 *ICEVchInd5* that was first described in a 2009 study characterizing a *V. cholerae* SXT ICE isolated from India in 1994^36^. These observations are consistent with the idea that a recent ancestor of ZTox was derived from *a V. cholerae* strain from South Asia. Other predicted genomic features of ZTox are identified in (S. Table 2)

### Generation of isogenic non-toxigenic Ogawa and Inaba derivatives of ZTox

We reasoned that deletion of *ctxAB* (CT) could potentially expand the utility of ZTox derivatives for use both as a challenge strain and as targets for vibriocidal assays, by removing the major diarrhoeagenic component, CT, of *V. cholerae*. The resulting strain, ZChol^O^ (Ogawa), had growth kinetics in standard laboratory conditions (LB media) that were indistinguishable from ZTox (Fig. S2A) and as expected did not produce CT when grown in laboratory ‘AKI’ conditions that induce CT synthesis in *V. cholerae* (Fig. S2B). The *wbeT* locus was deleted from ZChol^O^ to yield ZChol^I^, an Inaba version of this non-toxigenic strain (Fig. S2C). We anticipate that the non-toxigenic isogenic ZChol^O^ ZChol^I^ pair should be valuable to gauge serotype-specific responses to cholera and cholera vaccines. Both strains are susceptible to antibiotics such as azithromycin, tetracycline, combinations of sulfamethoxazole and trimethoprim, and ciprofloxacin (S. Table 1) that are often used to treat cholera; these agents could be used if these ZChol strains prove diarrheagenic in human studies.

### ZChol^O^ and ZChol^I^ colonize the infant mouse intestine but do not cause morbidity

We used the infant mouse model of cholera to compare the lethality as well as the intestinal colonization of ZChol^O^ and ZChol^I^ with the toxigenic ZTox parent strain. In this model, diarrhea and morbidity are dependent on CT activity^37^. A 10^7^ CFU oral dose routinely leads to diarrheal disease and death/moribundity within 48 hours with toxigenic *V. cholerae* strains^28,37,38^. All pups given ZTox developed diarrhea and became moribund by 35 hours post inoculation with a median time to death of 31 hours across 3 independent litters (Fig. 2A). In marked contrast, none of the animals inoculated with ZChol strains developed diarrhea or illness and all 6/6 and 6/6 of ZChol^O^ and ZChol^I^ respectively were alive 48 post-inoculation (Fig. 2A). Small intestinal CFU burdens were determined when animals became moribund (in the case of ZTox) or at 48 hours for ZChol strains. Despite the pronounced differences in the morbidity elicited by these strains, they all colonized the small intestine to a similar extent (Fig. 2B).

**Figure 2.**
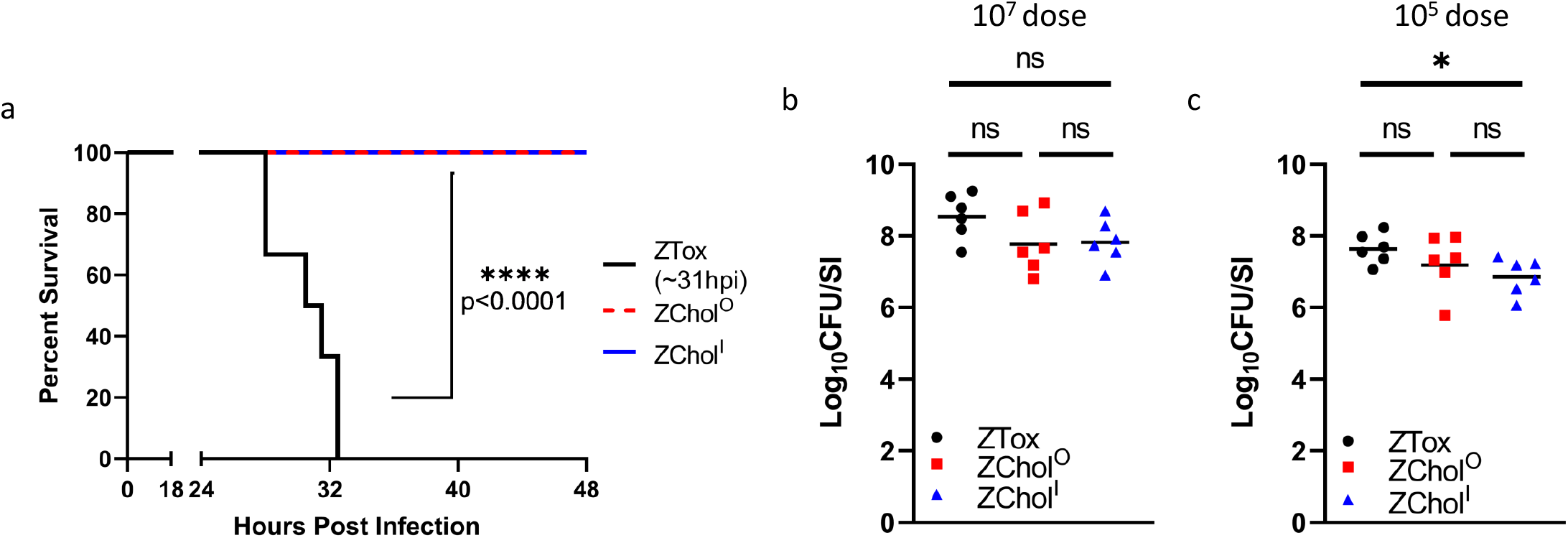
ZChol^O^ and ZChol^I^ robustly colonize the small intestine but do not cause death. **A)** Kaplan-Meier plot of P3 infant mice orally infected with a lethal dose of 10^7^ CFU of the indicated strains, n=6 per group. Median time to moribundity was 31 hours post infection for infant mice infected with the toxigenic parent strain. Differences in the survival curves were assessed with a log rank (Mantel Cox) test. **B)** *V. cholerae* burden in the small intestine (SI) of challenged mice at time of moribundity (ZTox) or at 48 hours post infection (ZChol strains). **C)** CFUs per SI for P5 infant mice orally infected with a non-lethal 10^5^ CFUs of indicated strains and sacrificed at 20 hours post inoculation. Differences in CFU burdens were assessed with a Mann-Whitney U test. Horizontal bars indicate geometric means of each group.

In a complementary experiment, a lower dose of 10^5^ CFU was used to directly compare the capacities of ZTox and ZChol^O^ and ZChol^I^ to colonize the SI at 20 hours post-inoculation, a point where less diarrheal disease was apparent. The small intestinal CFU burdens in infected mice were also similar using this protocol, though the ZChol^I^ burden was modestly lower than that of ZTox (Fig. 2C). Together, these data suggest that the engineered ZChol strains pose reduced risk of inducing cholera-like disease in human challenge studies, but nevertheless would be useful indicators of *V. cholerae* intestinal colonization.

### Utility of ZChol strains as challenge strains for vaccine studies

To model how ZChol strains could be used in human challenge studies, we measured how colonization with these strains could be used to determine vaccine efficacy (Fig. 3A). We immunized adult germ-free (GF) mice with killed and live formulations of the PanChol oral cholera vaccine^28,39^ and then challenged the pups of the immunized dams with either ZTox, ZChol^O^, or ZChol^I^ (Fig. 3A). In the GF model, live oral cholera vaccines are far more potent immunogens than killed whole cell cholera vaccines, which have minimal efficacy in this model as single dose agents^28^. Half of the pups from dams immunized with the killed vaccine exhibited diarrhea and inactivity upon ZTox challenge, whereas none of the 34 mice challenged with ZChol^O^ or ZChol^I^ developed signs of disease (Fig. 3B). Independent of challenge strain, the neonatal mice born to and suckled by dams that had received the live vaccine had significantly lower SI *V. cholerae* burdens than the mice from the killed vaccine group. This observation reflects the greater potency of the live vaccine and suggests that ZChol colonization can be used to gauge vaccine protection while reducing the risk inherent in challenge with toxigenic *V. cholerae*. In the killed PanChol vaccine group, the intestinal burden of ZChol strains was modestly lower in animals challenged with these non-toxigenic strains compared to ZTox (Fig. 3B). However, there were no significant differences in intestinal burdens of ZTox, ZChol^O^, or ZChol^I^ in animals in the live vaccine group. Together these data demonstrate the potential for ZChol strains to be used in de-risked human CHIM studies.

**Figure 3.**
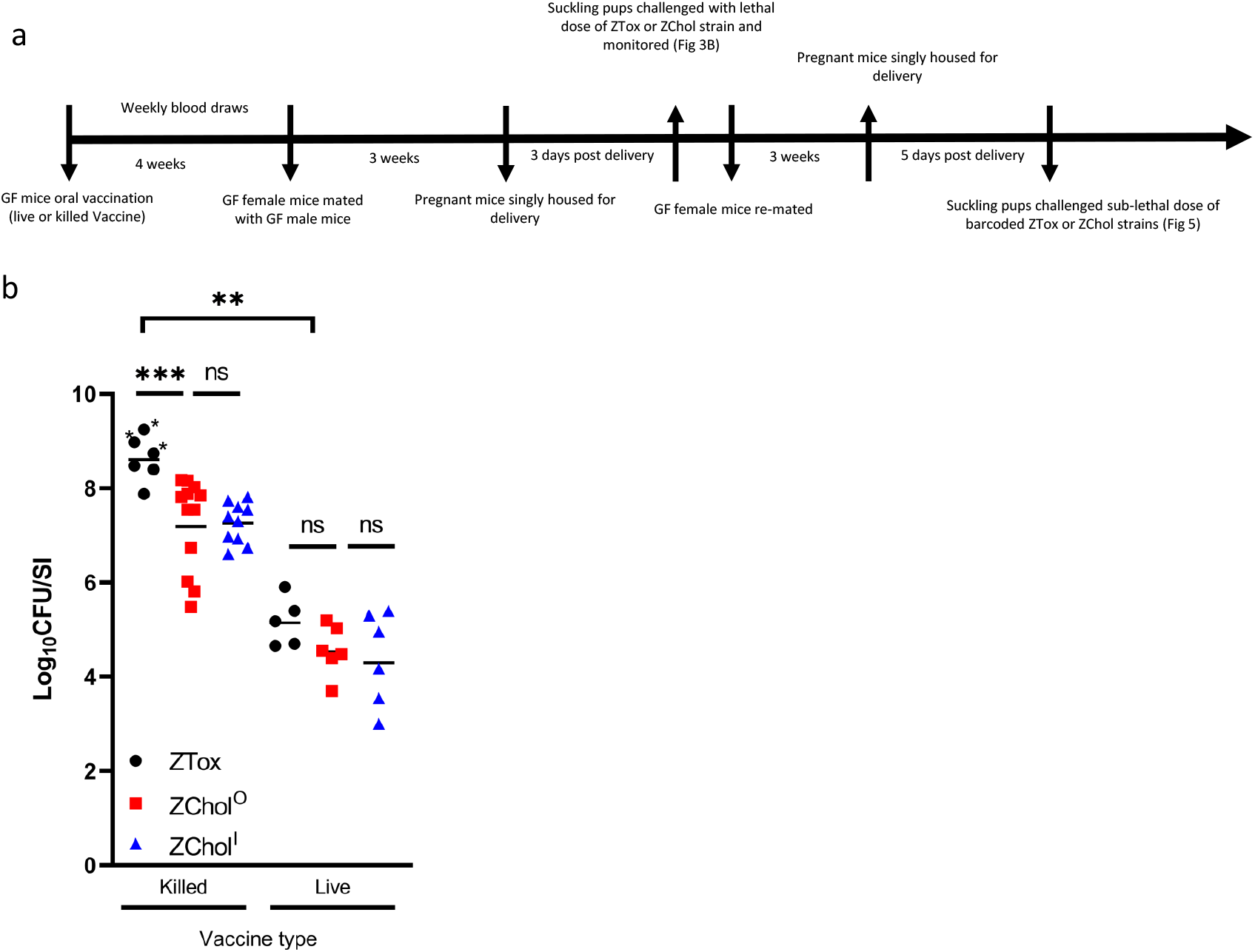
ZChol^O^ and ZChol^I^ provide a similar gauge of vaccine mediated protection as ZTox without causing diarrheal disease. **A)** Schematic overview of GF vaccination regimen and experimental protocol. **B)** P5 pups born to germ-free female mice orally immunized with either a live (n=5 GF dams) or killed oral cholera vaccine (OCV) (n=5 GF dams) were orally inoculated with ZTox or ZChol^O^ or ZChol^I^. CFU per SI in these pups were determined at 48 hours post inoculation (10^5^ CFU per infant mouse inoculum) with the indicated strains. Asterisks denote mice with visible diarrhea. Differences in CFU burdens were determined by a Mann-Whitney U test. Horizontal bars indicate geometric means of each group.

### ZChol strains are advantageous reagents for measuring vibriocidal antibody titers

The closest correlate of protective immunity against cholera are circulating vibriocidal antibody titers (VATs). Vibriocidal assays use Ogawa and Inaba strains, such as PIC158 and PIC018 respectively^40^, which are toxigenic and non-isogenic, as targets along with exogenous complement to measure the titer of complement fixing anti-Ogawa and anti-Inaba antibodies in serum samples. PIC018 was isolated from Bangladesh in 2007 and is relatively genetically distant from many currently circulating strains (Fig. 1); thus, using a more contemporary target strain may allow for greater sensitivity to contemporary *V. cholerae* antigens. We reasoned that ZChol strains, which are genetically identical except for the *wbeT* deletion, could be useful as target strains for measurements of serotype-specific vibriocidal antibodies, because they only differ in the methylation of the *V. cholerae* O1 O-antigen and not in other potential targets of vibriocidal antibodies. Furthermore, these strains pose lower biosafety risk to laboratory workers because they lack the CT genes.

We tested the ZChol pair as target strains in vibriocidal assays using murine serum obtained from a previous cholera vaccine study^28^, with the identical protocol used to measure serotype specific VATs in the previous study where PIC018 and PIC158 were used as the target strains. The correlation (r^2^ value) between the VATs reported using the PIC strains and those determined using the ZChol strains was 0.87 (Fig. 4A). In one sample, there was a detectable anti-Inaba VAT using ZChol^I^ that was not detected by the Inaba PIC018 strain (Fig. 4A), suggesting that ZChol^I^ may be a more sensitive indicator strain than PIC018.

**Figure 4.**
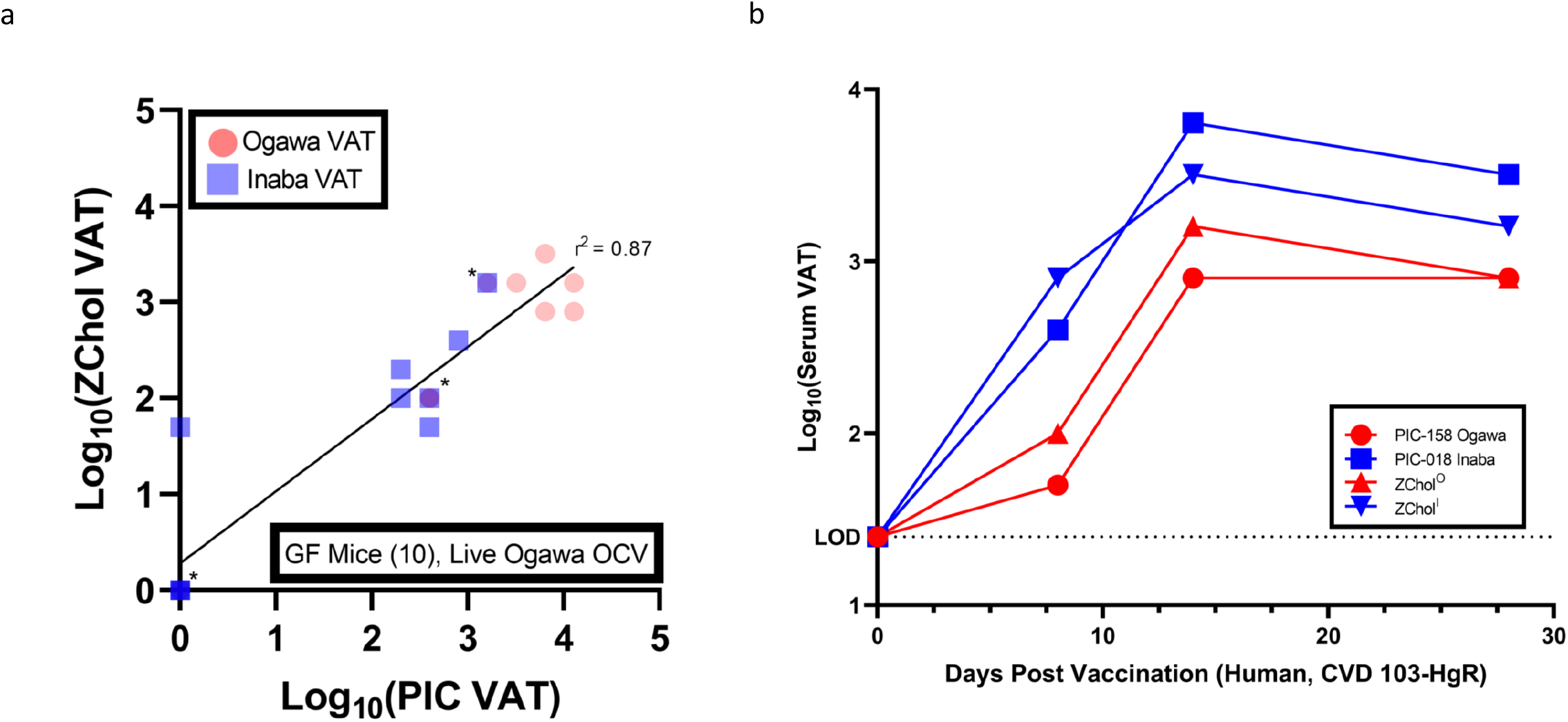
Measurement of vibriocidal antibody titers in both murine and human convalescent serum using ZChol^O^ or ZChol^I^ as targets. **A)** ZChol strains were used to determine the vibriocidal antibody titers (VATs) of live Ogawa OCV immunized GF mouse serum (n=10) from a previous study^28^ and compared to previously reported VAT values from non-isogenic PIC reporter strains. Red circles represent VATs determined by ZChol^O^ and PIC-158 (Ogawa) reporter strains, and the blue squares represent VATs determined by ZChol^I^ and PIC-018 (Inaba) reporter strains. VATs below the limit of detection were set to 1 for statistical analysis. The r^2^ linear correlation value between the PIC and ZChol strains was 0.87. Asterisks denote overlapping data points. **B)** ZChol strains and PIC strains were both used to determine the VATs from human serum obtained pre and post vaccination with CVD 103-HgR (Inaba live OCV). Dashed line represents the limit of detection (LOD) of this assay.

Similar experiments were carried out to quantify VATs from serum obtained from a human volunteer vaccinated with CVD103-HgR, a live-attenuated Inaba OCV. Again, ZChol strains, PIC158, and PIC018 were used as targets in parallel VAT assays (Fig. 3B). Both sets of reporter strains revealed the kinetics of the vibriocidal responses to a single dose of this live-attenuated vaccine, as well as the greater titer of anti-Inaba responses, which peaked 14 days post vaccination (Fig. 3B). We conclude that the ZChol matched strains are useful reagents for safely and accurately evaluating protection against cholera in this most widely used clinical assay.

### Barcoded challenge strains enable new insights into the mechanisms of vaccine protection

Reflecting the greater potency of live OCVs, mouse pups nursed by GF dams vaccinated with the live OCV had much lower intestinal *V. cholerae* burdens than those vaccinated with the killed OCV^28^. Two mechanisms could account for the greater passive protection (presumably mediated by maternal antibodies^37^) observed in the live OCV group. This reduction in *V. cholerae* burden could be explained by antibodies restricting the number of bacteria capable of initiating colonization (the founding population) and/or by impeding bacterial expansion, which represents bacterial replication minus death and loss through defecation. We introduced ~60,000 genomic barcodes into the genomes of ZTox, ZChol^O^, and ZChol^I^ and used our STAMPR analytic framework to distinguish between these possibilities^41^. STAMPR quantifies the founding population in a metric, Ns, that reflects the number of unique cells from an inoculum that give rise to the observed population in the host at the time of sampling. If passive immunity reduces bacterial expansion in the suckling pups, we expect Ns to remain unchanged while overall bacterial burden decreases. In contrast, if vaccination tightens host bottlenecks (thereby reducing the founding population size), we expect a decrease in Ns of the same magnitude as the decrease in CFU (Fig. 5A, model 2). If vaccination reduces both the founding population and bacterial expansion, we expect a decrease in Ns that is smaller in magnitude than the decrease in CFU (Fig. 5A, model 4).

**Figure 5.**
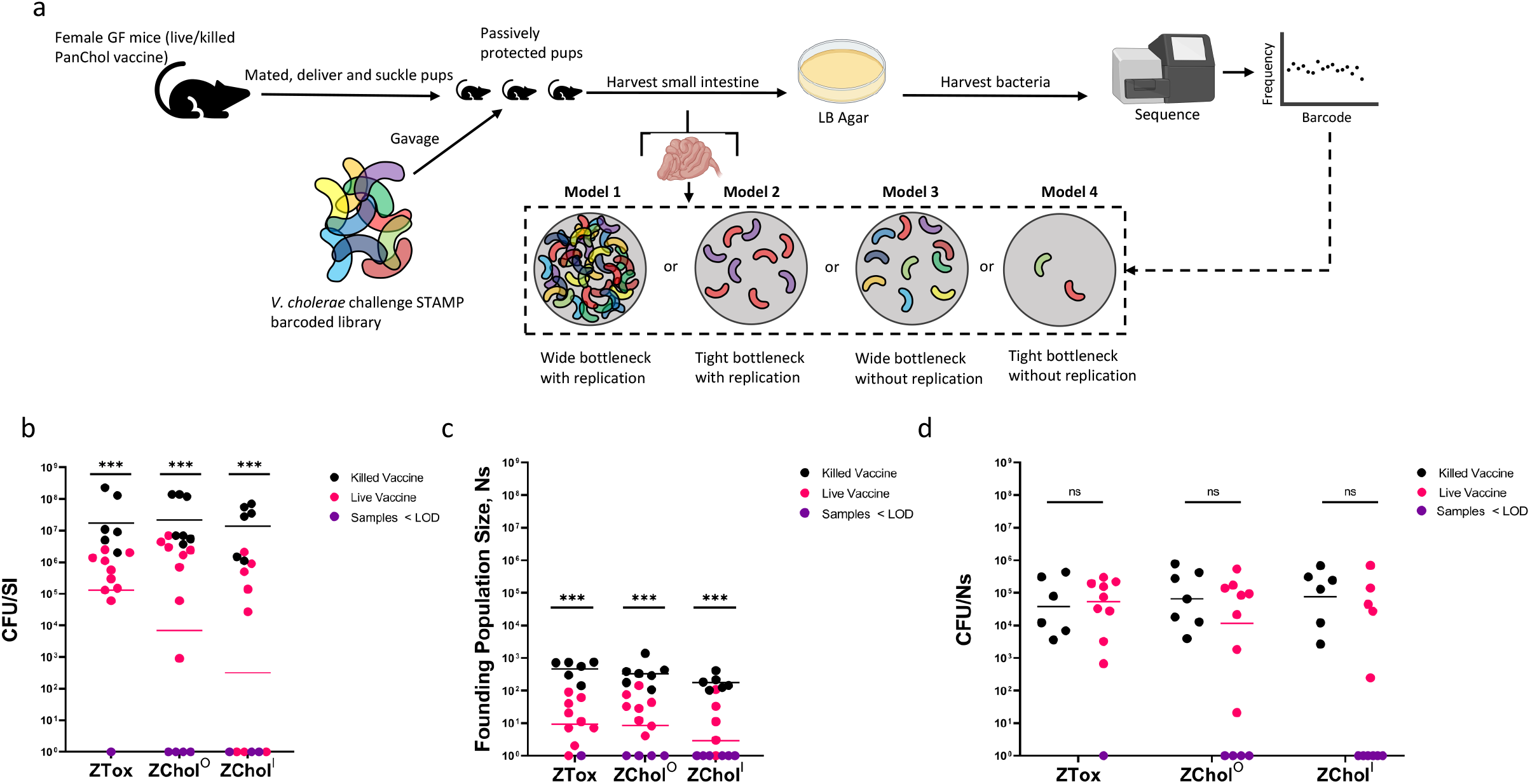
Barcoded ZChol strains can reveal the impact of vaccination on *V. cholerae* population dynamics during infection. **A)** Schematic of experimental protocol as well as interpretations of the potential outcomes of the barcode-based analyses of bacterial population composition. Infant mice from live or killed OCV vaccinated dams were challenged with genetically barcoded ZTox or ZChol^O^ or ZChol^I^ at a 10^5^ dose. Mice were sacrificed 20 hours post inoculation. **B)** CFU per SI from each mouse orally inoculated with the barcoded libraries. Purple circles are CFU per SI values below the limit of detection of this assay i.e., no recoverable CFUs from the inoculated infant mice. **C)** Ns values indicate the size of the *V. cholerae* founding populations, i.e., the number of bacteria from the 10^5^ CFU inoculum that successfully initiated colonization and expansion. Purple circles are Ns values below the limit of detection from deep sequencing. **D)** CFU/Ns values determined by the dividing the values in B by the values in C. Purple circles are CFU/Ns imputed from values below the limit of detection. Statistical differences were determined by Mann-Whitney U tests. Color-coded horizontal bars indicate geometric means of each group

As a control, Ns was first calculated across a wide range of known artificially created bottlenecks (Fig. S2D), confirming that this metric accurately quantifies bottlenecks for these libraries across 5 orders of magnitude. Then, P5 suckling infant mice from GF dams vaccinated with either the killed or live OCV as above were orally inoculated with the barcoded libraries of ZTox, ZChol^O^, or ZChol^I^ (Fig. 5A). As observed above (Fig. 3B), the burden of *V. cholerae* in pups nursed by dams vaccinated with the live vaccine were lower than those nursed by dams vaccinated with the killed OCV (Fig. 5B). The lower magnitude of difference in CFU per SI between the live and killed OCV groups (~1.5 log_10_ fold reduction) compared to those observed above (Fig. 3B) is consistent with waning immunity in the vaccinated dams. Independent of the challenge strain, the live vaccine uniformly restricted the Ns values detected compared to the killed PanChol OCV (Fig. 5C). These data are consistent with the model (Fig. 5A, model 2) that passively transferred antibodies elicited by OCVs restrict the pathogen’s bottleneck but has little impact on bacterial expansion in the host. Indeed, there is no significant difference in the CFU/Ns ratio (a gauge of expansion) between pups nursed by live PanChol OCV vaccinated or killed PanChol OCV vaccinated animals (Fig. 5D). Together, these findings reveal that antibodies elicited by vaccination almost exclusively influence host bottlenecks and suggest that maternal antibodies within milk limit the number of bacterial cells capable of initiating infection in the suckled infant mice. However, these antibodies appear to have little influence on the capacity of the *V. cholerae* cells that have passed through the bottleneck and reached suitable intestinal niches to expand. Thus, vaccination in this model creates a more restrictive bottleneck without impacting the expansion of the founding population.

## Discussion

Knowledge of *V. cholerae* pathogenesis and immunity has greatly benefitted from controlled human infection studies (CHIM) and CHIMs using toxigenic *V. cholerae* challenge strains have been instrumental for testing the protective efficacy of several cholera vaccine formulations. Here, we created ZChol^O^ and ZChol^I^, isogenic Ogawa and Inaba strains, respectively, as potential next generation *V. cholerae* challenge strains. These ZChol strains, which are derived from a contemporary *V. cholerae* isolate from Zambia, were rendered non-toxigenic to ameliorate safety concerns arising from CT-induced diarrhea. Potentially, these non-toxigenic strains might be used in outpatient studies, including in endemic countries, without the need for special facilities as are needed with wild type virulent challenge strains. Finally, thousands of barcodes were introduced into a neutral genetic locus in these strains, adding the potential for analyses of *V. cholerae* population dynamics to CHIM studies. Our pre-clinical analyses presented here suggest that these strains have the potential to enhance the safety, relevance, and scope of future CHIM studies.

We chose a recent Zambian *V. cholerae* isolate to create the ZChol Ogawa and Inaba pair because the highest burden of reported cholera cases is now in Sub-Saharan Africa. The strains have a set of genomic features that distinguish them from wave 1 strains such N16961 which is commonly used for CHIM studies. Genomic analyses (Fig. 1) show that ZTox is representative of modern pandemic cholera, including the strains that have given rise to extensive cholera outbreaks in the past 10-12 years in Haiti, Yemen, and multiple African countries^32–34^

Immune responses to CT are not the principal target of long-lasting protective immunity to cholera^42^. Instead, antibody responses targeting the O1 O-antigen (OSP) strongly correlate with protective immunity to cholera. Such antibodies, which can be elicited by infection or cholera vaccines, prevent *V. cholerae* colonization and lead to marked reductions in *V. cholerae* shedding in CHIM studies of immunity to cholera^24^. Given the reductions in *V. cholerae* shedding in stool in the setting of protective immunity, we reasoned that creation of non-toxigenic challenge strains may create a path to markedly de-risk future CHIM challenge studies. In the infant mouse model, the ZChol strains colonized the small intestine at comparable levels to ZTox, but in contrast to the latter strain, were well-tolerated (Fig. 2B). Furthermore, using a GF mouse model of vaccination and infant mouse challenge, we found that challenge with ZChol strains provided a similar measure of vaccine-elicited reduction of colonization of the challenge strain compared to ZTox (Fig. 3B). Since deletion of *ctxAB* does not always eliminate the diarrheagenicity of *V. cholerae* strains ^43^ it will be important to test whether the ZChol strains are minimally reactogenic prior to their adoption for human challenge studies.

While successful vaccination reduces the capacity of *V. cholerae* to colonize the intestine, the mechanisms that account for such reductions are not well-understood. We hypothesized that vaccine-associated reductions in *V. cholerae* colonization could be explained by immune response-mediated narrowing of the host bottleneck(s) to infection, limiting the size of the *V. cholerae* founding population and/or by immunity blocking of intestinal *V. cholerae* expansion (Fig. 5A, models). To distinguish between these possibilities, ZChol strains were barcoded with ~60,000 genomic sequence tags and we used STAMPR^41^ to quantify and compare the sizes of ZChol FPs in vaccinated animal models. Surprisingly, vaccination appeared to only narrow the host bottleneck to infection, reducing the number of *successful V. cholerae* founders, but did little to alter the ability of these founders to expand in the intestine. Thus, vaccine-elicited protective antibodies, which are thought to account for protection in this model, appear to largely limit the ability of *V. cholerae* to occupy its replicative niche in the small intestine. It will be highly informative to carry out similar experiments in the volunteer CHIM setting.

We propose that ZChol strains have the potential to dramatically alter the landscape of cholera CHIM studies and enable outpatient vaccine challenge studies in cholera endemic countries. Such studies could address pressing translational questions in the appropriate epidemiological settings, including optimization of dosing regimens for killed whole-cell vaccines like Shancol and Euvichol^44^ and testing of new cholera vaccine candidates such as HillChol^45^, PanChol^28^, and conjugate vaccines^46^, at far lower costs in critical settings in Africa and Asia. Beyond optimization of cholera vaccines, these new barcoded non-toxigenic strains should be valuable for studies of *V. cholerae* population dynamics in vaccinated and unvaccinated people across a broad set of circumstances, including studies of infectious doses in diverse settings where recipient nutritional status, prior exposure to the pathogen, and microbiota might influence mechanisms of immunity^47–49^. Finally, ZChol strains should also be valuable as safe and otherwise isogenic target strains for vibriocidal and immunologic assays of the host response to cholera.

## Methods

### Strains and growth conditions

A *V. cholerae* O1 2016 clinical isolate from Zambia (EDVRU/ZM/2016), named here ZTox, was serotyped by slide agglutination using Ogawa and Inaba specific antisera (BD Difco). Unless otherwise indicated all *V. cholerae* and *E. coli* strains were grown in LB medium and stored in 25% glycerol v/v at −80°C. For optimal induction of CT, *V. cholerae* was grown in AKI broth (0.5% sodium chloride, 0.3% sodium bicarbonate, 0.4% yeast extract, 1.5% Bacto peptone) for 4h without shaking, followed by 4h with aeration at 37°C^50,51^.

### Whole genome sequencing and phylogenetic analysis

Genomic DNA was extracted from ZTox with the Genejet Genomic DNA purification kit (Fisher) and prepared for whole-genome sequencing. DNA for short reads were prepared using the Illumina Nextera kit (Illumina) and sequenced using a NextSeq 550 (Illumina) at the Microbial Genome Sequencing Center (MIGS, Pittsburgh, PA). Long reads were generated using the Nanopore platform (Oxford) at MIGS. Adapter trimming and quality control standards were performed with bcl2fastq and a hybrid genome assembly was generated with Unicycler. The closed genome was then annotated using Prokka (MIGS, Pittsburgh). Raw fasta files were uploaded to Vicpred for further analyses^52^.

Raw sequence files and assembled genomes were analyzed as in^31^. Briefly, raw sequence files were downloaded from the European Nucleotide Archive, while 100 bp overlapping simulated reads were generated from the assembled genomes by using fasta_to_fastq.pl (https://github.com/ekg/fastato-fastq/blob/master/fasta_to_fastq.pl) or wgsim (https://github.com/lh3/wgsim). Simulated paired-end reads and paired-end reads were then mapped to the reference genome of 7PET wave 1 isolate N16961 (Genbank accession LT907989 and LT907990) with Snippy v4.6.0 (https://github.com/tseemann/snippy) using freebayes v1.3.5 (https://github.com/freebayes/freebayes) with requirements for a mapping quality of 60, minimum base quality of 13, minimum read coverage of 4, and a 75% read concordance at a locus required for reporting. Alignments from^31^ were kindly provided by D. Domman (University of New Mexico). The recombinogenic VSP-II region, as well as the repetitive TLC-RS1-CTX region were then masked from the alignment using Geneious (Biomatters). Further recombinogenic sites were masked using Gubbins v3.1.2^53^. A maximum-likelihood phylogenetic tree was generated from ~10,800 genomic SNPs using RAxML v.8.2.12 ^54^ with the GTR model and 100 bootstraps. The tree was rooted on the pre-seventh pandemic A6 genome and visualized in iTOL v5^55^.

The SXT flanking sequence “ATCATCTCGCACCCTGA’’ was chosen from a previous comparative SXT ICE study ^56^ and identified in the ZTox genome. The other flank of the SXT region was then defined as the 5’ end of the *prfC* gene ^57^. The resulting ZTox-SXT was identified and aligned to the H1-SXT ^58^ from in Geneious (Biomatters) using MUSCLE alignment.

### Creation of ZChol^O^ and ZChol^I^ strains

A streptomycin resistant variant of ZTox was isolated by plating a concentrated overnight ZTox culture onto streptomycin (Sm) LB agar plates (1 ng/mL) and grown for 24 hours at 30°C. A streptomycin resistant (SmR) colony was picked, grown in LB Sm (200 μg/mL), and frozen as a glycerol stock. Whole genome sequencing identified the resistance mutation as *rpsL* A263G (K88R), a common allele conferring resistance to Sm^59^.

A nontoxigenic (*ΔctxAB*) derivative of ZTox, ZChol^O^, was generated by allelic exchange. Initially, pBF31, a derivative of the allele exchange vector pCVD442 (Carbenicillin resistance, Carb) containing the kanamycin (Kan) resistance cassette from pKD4^60^ flanked by FRT sites sandwiched by 1kb homology arms targeting the *ctxAB* operon was created. BF31 was conjugated from SM10λpir *E. coli* into SmR ZTox and single crossover mutants were selected on LB+Sm200μg/Carb50μg/Kan50μg/mL agar plates. Colonies were then grown in liquid LB+Carb50/Kan50 at 37°C for 6 hours and then inoculated into static LB+10% sucrose for 24 hours at room temperature. Sucrose resistant (sacB-negative), Carb^S^, and Kan^R^ colonies were isolated and checked by colony PCR using internal and flanking primers to demonstrate the loss of *ctxAB* and the insertion of the FRT flanked kanamycin resistance cassette. The FRT flanked Kan^R^ cassette was removed by introducing pCP20^60^, which encodes the Flp recombinase. To accomplish this, a single colony of ZToxΔ*ctxAB*-FRT-Kan^R^ was grown in 50mL LB with Sm200+Kan50 μg/mL ON at 37°C shaking at 220rpm and then pelleted. Pellets were washed 2X with ice-cold water and then 2X with ice-cold 10% glycerol to render them electrocompetent. 50μL of electrocompetent ZToxΔ*ctxAB*-FRT-Kan^R^ were electro-transformed with 50ng of pCP20, recovered in 1mL SOB for 3 hours, and then plated on LB agar+Sm200+Carb50 and grown ON at 30°C. Individual Sm^R^Carb^R^ colonies were inoculated into LB and grown at 42°C for 10 hours and then serially diluted onto LB agar+Sm200 ON at 37°C. Individual colonies were replica struck onto LB agar+Sm200μg/mL, LB Sm200/Kan50μg/mL agar, and LB Sm200/Kan50/Carb50μg/mL agar. Sm^R^Kan^S^Carb^S^ colonies were tested by colony PCR to verify the loss of the FRT-Kan^R^ yielding *ZToxΔctxAB* (ZChol^O^). The growth of ZTox and ZChol^O^ in LB were compared in a 96 well plate format and OD_600_ was determined by a plate reader to generate growth curves.

ZChol^O^ was mated with SM10λpir *E. coli* carrying pBF32 (pCVD442-WbeT), which contains 1 kb homology arms targeting the *wbeT* locus, and allelic exchange was carried out as described above using pCVD442 and *sacB* counterselection to isolate *ΔwbeT* strains. Colony PCR was used to confirm the resultant strain was ZChol^O^Δ*wbeT* (ZChol^I^). Short read whole genome sequencing (Illumina, MIGS) was performed on both ZChol strains to verify the intended mutations. Slide agglutination with *V. cholerae* serotype specific antisera (BD Difco) was performed using glass slides to confirm serotypes. Antibiotic susceptibility assays were performed using E-test strips (Biomeriux) and the results interpreted using the CLSI M45-A3 2016 guidelines.

### Western blot analysis

Proteins were separated by SDS-PAGE using 4-12% NuPAGE Bis/Tris precast gels (Life Technologies) and transferred to nitrocellulose using an iBlot gel transfer device (Life Technologies). The pre-stained protein marker, SeeBlue (Invitrogen) was used as molecular mass standards. Rabbit anti-CT polyclonal antibody (Abcam ab123129) and horseradish peroxidase (HRP)-linked anti-rabbit IgG were used as primary and secondary antibodies, respectively. Blots were developed with the SuperSignal West Pico PLUS Chemiluminescence substrate (ThermoFisher) and exposed in a Chemidoc system (Bio-Rad Laboratories).

### Infant mouse lethal dose challenge assay

Infant mouse lethal dose challenges were performed as previously described^37,38^. Five female C57BL/6 dams with 2–3-day old litters (P2-3) (Charles River) were singly housed with their pups. At P3-4, the pups within each litter were randomly assigned to three groups, orally inoculated with 10^7^ CFU in 50μL LB of the indicated *V. cholerae* strain and returned to their dams for maternal care. Inocula were serially diluted and plated in triplicate for CFU enumeration to confirm infection dose. Infected pups were closely monitored every 2-4 hours for onset of diarrhea and diminished activity levels. At this point, monitoring was increased to 30-minute intervals until pups reached moribundity. When pups became moribund or at the 48-hour time-point, pups were euthanized for necropsy, SI weighing, homogenization, and plating on LB Sm200 agar to enumerate the CFU per SI of the challenged mice was as described previously^28,37^.

### Infant mouse colonization assay

Three litters of 4–5-day old C57BL/6 infant mice (Charles River) were separated from their dams upon receipt and placed into isolation incubators with padding and nesting material. Inocula were prepared by making 1:1000 dilutions of 37°C overnight LB cultures of the indicated strains (roughly 10^5^ CFU/50 μL inocula). The infant mice were randomly split into three groups and each mouse was orally inoculated with the indicated strain and returned to the incubation chambers for 20 hours. After 20 hours, the mice were euthanized and necropsied and CFU/SI, were enumerated as described above.

### Germ-free immunization study

Ten 4-week-old C57BL/6 female germ-free (GF) mice (Massachusetts Host-Microbiome Center) were maintained in a BL2 facility with autoclave-sterilized cages, food, and water. On day 0 of the experiment, half (five) of the mice were anesthetized and orally inoculated with 10^9^ CFU of an overnight culture of live PanChol oral cholera vaccine (OCV) in 100μL sodium bicarbonate^28^. The other half of the mice were anesthetized and given the same dose (10^9^ CFU) of formalin-killed PanChol in sodium bicarbonate (Fig. 3A). Mice were co-housed according to their vaccine group. 4 weeks post-vaccination, the female mice were mated with age matched GF C57BL/6 male mice and monitored closely for signs of pregnancy. Pregnant dams were singly housed in sterilized cages beginning at ~E-18 (three days before delivery) for delivery. Randomly assigned suckling P3 pups from each litter were challenged with typically lethal doses (10^7^ CFU) of the indicated strains as described and returned to their respective dams for care. CFU per SI enumeration after 48 hours of infection of the pups was carried out as previously described.

### Vibriocidal antibody titers assays

Murine and human vibriocidal antibodies were quantified by finding the highest serum dilution required to lyse isogenic ZChol strains or PIC158 (Ogawa) or PIC018 (Inaba) *V. cholerae* as described previously^28,61^. Heat inactivated 2-fold serial dilutions of serum were incubated with guinea pig complement (Sigma) and the target strain, and then allowed to grow in brain heart infusion (BHI) medium in a 96-well plate. The serum dilution that caused more than 50% reduction in target strain OD_595_ compared to saline negative control wells was recorded as the antibody titer. A mouse monoclonal antibody, 432A.1G8.G1.H12^28^, targeting *V. cholerae* O1 OSP was a positive control for the assay. The limit of detection represents the lowest serum dilution at which no inhibition of growth could be detected.

### Barcode analysis of population bottlenecks

To create a donor plasmid containing ~60,000 barcodes, a fragment containing the Tn7 transposon and necessary conjugation/replication machinery from pJMP1339^62^ was generated. In addition, we amplified a kanamycin resistance cassette from pDS132-STAMPR ^41^ with primers containing 25 ‘N’ nucleotides, representing ~1 x 10^15^ possible barcodes. These fragments were assembled with the NEB HiFi DNA Master Mix to create pSM1. The assembly was then electroporated at scale into MFDλpir and ~60,000 colonies were pooled and frozen into several aliquots.

To introduce barcodes into recipient *V. cholerae* strains, 30μL of thawed donor culture (MFDλpir+pSM1) was inoculated in 3mL of LB Kan50/Carb50/DAP300μg/mL, grown overnight and mixed with overnight cultures of recipient *V. cholerae* strains at equal ratios (total volume of 1.2mL) along with the donor containing the helper transposase (MFDλpir+pJMP1039). Cells were pelleted, resuspended in 100μL of LB and spotted on a 0.45μm HAWP filter (Millipore). This procedure was scaled 6x to create the final library. After incubation for 4 hours at 37°C, cells were washed from the filter and plated on LB Kan50μg/mL agar. The absence of DAP counterselects against the donor strain. The resulting transconjugant colonies were then pooled and frozen and used for future infant mouse inoculations.

STAMPR library preparation was performed as previously described^41^. Briefly, 2ul of bacterial suspension was diluted in 100ul of water and boiled for 15 min at 95°C. PCR to amplify the barcode region was performed in 2ul of boiled cells using Phusion DNA polymerase (New England Biolabs). The presence of the correct PCR product was verified on an agarose gel and samples were pooled, purified, and sequenced on a MiSeq (Illumina) for 78 cycles. Sequence analysis was performed using the CLC genomics workbench (Qiagen). To create the reference list of barcodes, we first sequenced the MFD donor library as done previously^41^. After trimming the reads, sequences were deduplicated in Geneious (Biomatters) using the dedupe plugin with default settings. This resulted in a preliminary list of ~100,000 barcodes (reference list RL1). We then deep sequenced the *V. cholerae* transconjugants and mapped the reads onto RL1. Consensus sequences for barcodes with at least one read were obtained to create RL2. The mapping procedure was repeated and consensus sequences for barcodes containing more than one read were collected, creating RL3. RL3 was deduplicated in Geneious as above to create RL4. Reads were then mapped again to RL4 and consensus sequences for barcodes containing more than 1 read were obtained, to create the final reference list containing 63209 barcodes.

Reads from all samples were mapped to this reference list of barcodes and exported as a table of read counts. From these counts, we calculated Ns as done previously ^41^, with several modifications to bolster the initial correction for sequencing noise. Since the number of barcodes (~60,000) is close to the sequencing depth (~100,000-200,00), many barcodes have only one read mapping to them. We define m as the number of barcodes with greater than 1 read mapping and n as the number of barcodes with only 1 read mapping. The m/n ratio is used to define a noise correction factor proportional to the sequencing depth (ss). A noise correction factor C is then defined as:

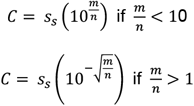

We then sampled the distribution of the entire sequencing run (which represents the noise distribution that arises by index hopping) via multinomial resampling C times and subtracted these reads from the output samples. One read was subtracted from all barcodes if m/n < 0.83 after this initial noise correction. In practice, these corrections remove reads from samples that are overrepresented by barcodes with only one read mapping to them, as these are likely to be noise. The remaining noise is identified by local minima in the Resiliency algorithm ^41^.

In addition, we also used the read depth of the output sample to simulate a bottleneck on the input, prior to Ns determination. This correction adjusts for the case when more barcodes would have been detected had the sample been sequenced more deeply. The full code used to analyze these data is provided in https://github.com/hullahalli/stampr_rtisan.

### Statistical analysis

Statistical analyses were performed with Prism 9 (GraphPad). Specific statistical tests used for analysis are defined in the Figure legends.

### Animal use statement

All experiments in this study were performed as approved by the Brigham and Women’s Hospital IACUC (protocol 2016N000416) and in compliance with the Guide for the Care and Use of Laboratory Animals.

## Acknowledgements

We acknowledge laboratory members in the Waldor, Sack and Chilengi labs, and Dr Michelo Simuyandi for their contributions and discussion and Dr. Domman for conversations and advice with phylogenies. We also thank the Massachusetts Host-Microbiome Center for assistance with mouse procurement and husbandry. Figure 5 was generated using www.biorender.com.

## Author Contributions

B.F., D.S., R.C., M.K.W. conceived the study. B.F. performed most of the experiments. K.H. created, sequenced, and analyzed the barcode libraries. D.H.F.R. performed the phylogenetic analyses. D.R.L. assisted with some animal work and phenotypical characterization. B.F. wrote the manuscript with input from all other authors. R.C., D.S., and M.K.W. provided scientific direction and M.K.W. allocated funding for this study.

## Supplementary Figures

**S1**

**A)** Genomic organization of ZChol CTXΦ prophage as well as the RS1 satellite phage. The cholera toxin genes are indicated in red and were deleted from ZTox to generate ZChol^O^.

**B)** Comparison of SXT ICEs found in ZTox and 7PET H1 *V. cholerae*. The annotated ZTox-SXT has almost complete consensus homology with the H1-SXT ICE (green consensus bar) by nucleotide alignment with MUSCLE. Like the *ICEVchInd5* originally described in a *V. cholerae* strain isolated in 1994 India^36^, the ZTox-SXT lacks a 12.49kb region that codes for multiple antibiotic resistance genes as well as two previously unannotated predicted proteins. The borders of the missing region are flanked by predicted transposases.

**S2**

**A)** Neither the *rpsL* K88R streptomycin resistance mutation nor deletion of *ctxAB* influences the growth rate of ZChol^O^ strains under standard laboratory conditions.

**B)** Western blot analysis of CT production under standard laboratory growth conditions as well as ‘AKI’ virulence gene induction conditions. Immunoblots were loaded with equal amounts of sterile-filtered supernatants of AKI grown ZTox (lane 1) and ZChol^O^ (lane 2) as well as with sterile-filtered supernatants of LB grown ZTox (lane 3) and ZChol^O^ (lane 4) and incubated with an anti-CT polyclonal antibody (Abcam ab123129). The sizes of the A and A1-subunit (A and A1) as well as the B-subunit (B) of CT are indicated by arrows on the right. Lines to the left indicate the molecular masses of the protein standards in kDa.

**C)** Slide agglutination analysis using serotype specific anti-serum demonstrates that ZChol^O^ and ZChol^I^ are Ogawa and Inaba serotypes, respectively.

**D)** Barcoded ZTox and ZChol strains were serially diluted and plated for CFU and then enumerated. Sequencing of these libraries showed that the Ns:CFU relationship correlated well and that Ns is a good predictor of known CFU values.

## Supplementary Tables

**Table 1)**Both ZChol^O^ and ZChol^I^ are sensitive to antibiotics commonly used to treat *V. cholerae*. S = sensitive, R = resistant, N = Indeterminate.

**Table 2)**Predicted gene contents of *V. cholerae* pathogenicity islands using the Vicpred^52^ analysis pipeline.

## References

1. Ali, M., Nelson, A. R., Lopez, A. L. & Sack, D. A. Updated global burden of cholera in endemic countries. PLoS Negl. Trop. Dis. 9, e0003832 (2015).

2. Troeger, C. et al. Estimates of the global, regional, and national morbidity, mortality, and aetiologies of diarrhoea in 195 countries: a systematic analysis for the Global Burden of Disease Study 2016. Lancet Infect. Dis. 18, 1211–1228 (2018).

3. Clemens, J. D., Nair, G. B., Ahmed, T., Qadri, F. & Holmgren, J. Cholera. Lancet (London, England) 390, 1539–1549 (2017).

4. Pezzoli, L. Global oral cholera vaccine use, 2013-2018. Vaccine 38 Suppl 1, A132–A140 (2020).

5. Shaikh, H., Lynch, J., Kim, J. & Excler, J. L. Current and future cholera vaccines. Vaccine 38, A118–A126 (2020).

6. GTFCC. Global Task Force on Cholera Control: Roadmap 2030. Global Task Force on Cholera Control https://www.gtfcc.org/about-gtfcc/roadmap-2030/ (2020).

7. Hisatsune, K., Kondo, S., Isshiki, Y., Iguchi, T. & Haishima, Y. Occurrence of 2-O-Methyl-N-(3- Deoxy-L-glycero-tetronyl)-D-perosamine (4-amino-4,6-dideoxy-D-manno-pyranose) in Lipopolysaccharide from Ogawa but Not from Inaba O Forms of O1 Vibrio cholerae. Biochem. Biophys. Res. Commun. 190, 302–307 (1993).

8. Devault, A. M. et al. Second-Pandemic Strain of Vibrio cholerae from the Philadelphia Cholera Outbreak of 1849. http://dx.doi.org/10.1056/NEJMoa1308663 370, 334–340 (2014).

9. Ramamurthy, T. et al. Revisiting the Global Epidemiology of Cholera in Conjunction With the Genomics of Vibrio cholerae. Front. Public Heal. 7, 203 (2019).

10. Mutreja, A. et al. Evidence for several waves of global transmission in the seventh cholera pandemic. Nature 477, 462–465 (2011).

11. Mengel, M. A., Delrieu, I., Heyerdahl, L. & Gessner, B. D. Cholera outbreaks in Africa. in Current Topics in Microbiology and Immunology vol. 379 (2014).

12. Sack, D. A. et al. Contrasting Epidemiology of Cholera in Bangladesh and Africa. J. Infect. Dis. (2021) doi:10.1093/INFDIS/JIAB440.

13. Levine, M. M. et al. Duration Of Infection-Derived Immunity To Cholera. J. Infect. Dis. 143, 818–820 (1981).

14. Cash, R. A. et al. Response of Man to Infection with Vibrio cholerae. I. Clinical, Serologic, and Bacteriologic Responses to a Known Inoculum. J. Infect. Dis. 129, 45–52 (1974).

15. Ali, M., Emch, M., Park, J. K., Yunus, M. & Clemens, J. Natural cholera infection-derived immunity in an endemic setting. J. Infect. Dis. 204, 912–918 (2011).

16. Holmgren, J., Clemens, J., Sack, D. A. & Svennerholm, A. M. New cholera vaccines. Vaccine 7, 94–96 (1989).

17. Ryan, E. T., Calderwood, S. B. & Qadri, F. Live attenuated oral cholera vaccines. Expert Review of Vaccines vol. 5 483–494 (2006).

18. Plotkin, S. A. & Gilbert, P. B. Nomenclature for immune correlates of protection after vaccination. Clin. Infect. Dis. 54, 1615–1617 (2012).

19. Holmgren, J. et al. Correlates of protection for enteric vaccines. Vaccine 35, 3355–3363 (2017).

20. Bhattacharya, S. K. et al. 5 year efficacy of a bivalent killed whole-cell oral cholera vaccine in Kolkata, India: a cluster-randomised, double-blind, placebo-controlled trial. Lancet. Infect. Dis. 13, 1050–1056 (2013).

21. Black, R. E. et al. Protective efficacy in humans of killed whole-vibrio oral cholera vaccine with and without the B subunit of cholera toxin. Infect. Immun. 55, 1116–1120 (1987).

22. Clemens, J. D. et al. Field trial of oral cholera vaccines in Bangladesh: results from three-year follow-up. Lancet (London, England) 335, 270–273 (1990).

23. Bi, Q. et al. Protection against cholera from killed whole-cell oral cholera vaccines: a systematic review and meta-analysis. Lancet Infect. Dis. 17, 1080–1088 (2017).

24. Chen, W. H. et al. Single-dose live oral cholera vaccine CVD 103-HgR protects against human experimental infection with vibrio cholerae O1 El Tor. Clin. Infect. Dis. 62, 1329–1335 (2016).

25. Cohen, M. B. et al. Randomized, controlled human challenge study of the safety, immunogenicity, and protective efficacy of a single dose of Peru-15, a live attenuated oral cholera vaccine. Infect. Immun. 70, 1965–1970 (2002).

26. Levine, M. M. et al. Volunteer studies of deletion mutants of Vibrio cholerae O1 prepared by recombinant techniques. Infect. Immun. 56, 161–167 (1988).

27. Erdem, R. et al. A Phase 2a randomized, single-center, double-blind, placebo-controlled study to evaluate the safety and preliminary efficacy of oral iOWH032 against cholera diarrhea in a controlled human infection model. PLoS Negl. Trop. Dis. 15, e0009969 (2021).

28. Sit, B., Fakoya, B., Zhang, T., Billings, G. & Waldor, M. K. Dissecting serotype-specific contributions to live oral cholera vaccine efficacy. Proc. Natl. Acad. Sci. 118, (2021).

29. Darton, T. C. et al. Design, recruitment, and microbiological considerations in human challenge studies. The Lancet Infectious Diseases vol. 15 (2015).

30. Gordon, S. B. et al. A framework for Controlled Human Infection Model (CHIM) studies in Malawi: Report of a Wellcome Trust workshop on CHIM in Low Income Countries held in Blantyre, Malawi. Wellcome Open Res. 2, (2017).

31. Mashe, T. et al. Highly Resistant Cholera Outbreak Strain in Zimbabwe. N. Engl. J. Med. 383, 687–689 (2020).

32. Weill, F. X. et al. Genomic history of the seventh pandemic of cholera in Africa. Science (80−.). 358, 785–789 (2017).

33. Weill, F. X. et al. Genomic insights into the 2016–2017 cholera epidemic in Yemen. Nat. 2019 5657738 565, 230–233 (2019).

34. Bwire, G. et al. Molecular characterization of Vibrio cholerae responsible for cholera epidemics in Uganda by PCR, MLVA and WGS. PLoS Negl. Trop. Dis. 12, (2018).

35. Davis, B. M., Kimsey, H. H., Kane, A. V. & Waldor, M. K. A satellite phage-encoded antirepressor induces repressor aggregation and cholera toxin gene transfer. EMBO J. 21, 4240 (2002).

36. Wozniak, R. A. F. et al. Comparative ICE Genomics: Insights into the Evolution of the SXT/R391 Family of ICEs. PLOS Genet. 5, e1000786 (2009).

37. Sit, B. et al. Oral immunization with a probiotic cholera vaccine induces broad protective immunity against vibrio cholerae colonization and disease in mice. PLoS Negl. Trop. Dis. 13, e0007417 (2019).

38. Fakoya, B., Sit, B. & Waldor, M. K. Transient intestinal colonization by a live-attenuated oral cholera vaccine induces protective immune responses in streptomycin-treated mice. J. Bacteriol. (2020) doi:10.1128/jb.00232-20.

39. Hubbard, T. P. et al. A live vaccine rapidly protects against cholera in an infant rabbit model. Sci. Transl. Med. 10, eaap8423 (2018).

40. Azman, A. S. et al. Estimating cholera incidence with cross-sectional serology. Sci. Transl. Med. 11, (2019).

41. Hullahalli, K., Pritchard, J. R. & Waldor, M. K. Refined Quantification of Infection Bottlenecks and Pathogen Dissemination with STAMPR. mSystems 6, (2021).

42. Levine, M. M. et al. Immunity of cholera in man: Relative role of antibacterial versus antitoxic immunity. Trans. R. Soc. Trop. Med. Hyg. 73, 3–9 (1979).

43. Rui, H. et al. Reactogenicity of live-attenuated Vibrio cholerae vaccines is dependent on flagellins. Proc. Natl. Acad. Sci. U. S. A. 107, 4359–4364 (2010).

44. Lopez, A. L. et al. Immunogenicity and Protection From a Single Dose of Internationally Available Killed Oral Cholera Vaccine: A Systematic Review and Metaanalysis. Clin. Infect. Dis. An Off. Publ. Infect. Dis. Soc. Am. 66, 1960 (2018).

45. Sharma, T. et al. Development of Hillchol^®^, a low-cost inactivated single strain Hikojima oral cholera vaccine. Vaccine 38, 7998–8009 (2020).

46. Sayeed, M. A. et al. A cholera conjugate vaccine containing ospecific polysaccharide (OSP) of V. cholera o1 inaba and recombinant fragment of tetanus toxin heavy chain (OSP:rTTHC) induces serum, memory and lamina proprial responses against OSP and is protective in mice. PLoS Negl. Trop. Dis. 9, e0003881 (2015).

47. Sack, G. H. et al. Gastric acidity in cholera and noncholera diarrhoea. Bull. World Health Organ. 47, 31 (1972).

48. Van Loon, F. P. L. et al. Low gastric acid as a rise factor for cholera transmission: Application of a new non-invasive gastric acid field test. J. Clin. Epidemiol. 43, 1361–1367 (1990).

49. Alavi, S. et al. Interpersonal Gut Microbiome Variation Drives Susceptibility and Resistance to Cholera Infection. Cell 181, 1533–1546.e13 (2020).

50. Iwanaga, M. & Yamamoto, K. New medium for the production of cholera toxin by Vibrio cholerae 01 biotype El Tor. J. Clin. Microbiol. 22, 405–408 (1985).

51. Iwanaga, M. et al. Culture Conditions for Stimulating Cholera Toxin Production by Vibrio cholerae 01 El Tor. Microbiol. Immunol. 30, 1075–1083 (1986).

52. Lee, I. et al. VicPred: A Vibrio cholerae Genotype Prediction Tool. Front. Microbiol. 12, (2021).

53. Croucher, N. J. et al. Rapid phylogenetic analysis of large samples of recombinant bacterial whole genome sequences using Gubbins. Nucleic Acids Res. 43, e15 (2015).

54. Stamatakis, A. RAxML-VI-HPC: maximum likelihood-based phylogenetic analyses with thousands of taxa and mixed models. Bioinformatics 22, 2688–2690 (2006).

55. Letunic, I. & Bork, P. Interactive Tree Of Life (iTOL) v5: an online tool for phylogenetic tree display and annotation. Nucleic Acids Res. 49, W293–W296 (2021).

56. Wang, R., Yu, D., Yue, J. & Kan, B. Variations in SXT elements in epidemic Vibrio cholerae O1 El Tor strains in China. Sci. Rep. 6, (2016).

57. Hochhut, B. & Waldor, M. K. Site-specific integration of the conjugal Vibrio cholerae SXT element into prfC. Mol. Microbiol. 32, 99–110 (1999).

58. Chin, C. S. et al. The origin of the Haitian cholera outbreak strain. N. Engl. J. Med. 364, 33–42 (2011).

59. Allué-Guardia, A., Echazarreta, M., Koenig, S. S. K., Klose, K. E. & Eppinger, M. Closed genome sequence of Vibrio cholerae O1 El Tor Inaba strain A1552. Genome Announc. 6, (2018).

60. Datsenko, K. A. & Wanner, B. L. One-step inactivation of chromosomal genes in Escherichia coli K- 12 using PCR products. Proc. Natl. Acad. Sci. U. S. A. 97, 6640 (2000).

61. Son, M. S. & Taylor, R. K. Vibriocidal assays to determine the antibody titer of patient sera samples. Curr. Protoc. Microbiol. 23, 6A.3.1–6A.3.9 (2011).

62. Peters, J. M. et al. Enabling genetic analysis of diverse bacteria with Mobile-CRISPRi. Nat. Microbiol. 2019 42 4, 244–250 (2019).

