## Supplementary figures and images for "Non-toxigenic *Vibrio cholerae* challenge strains for evaluating vaccine efficacy and inferring mechanisms of protection"

### Supplemental Figures

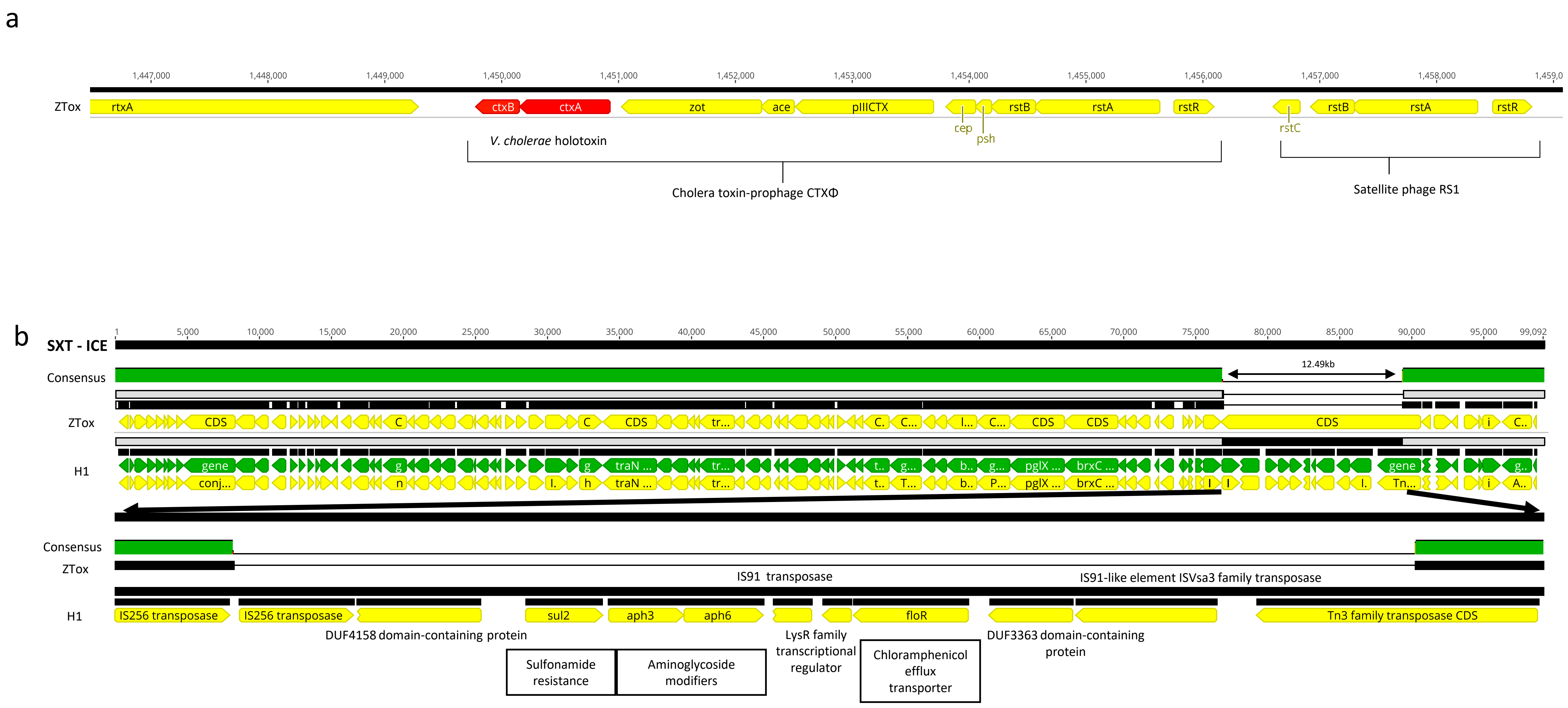

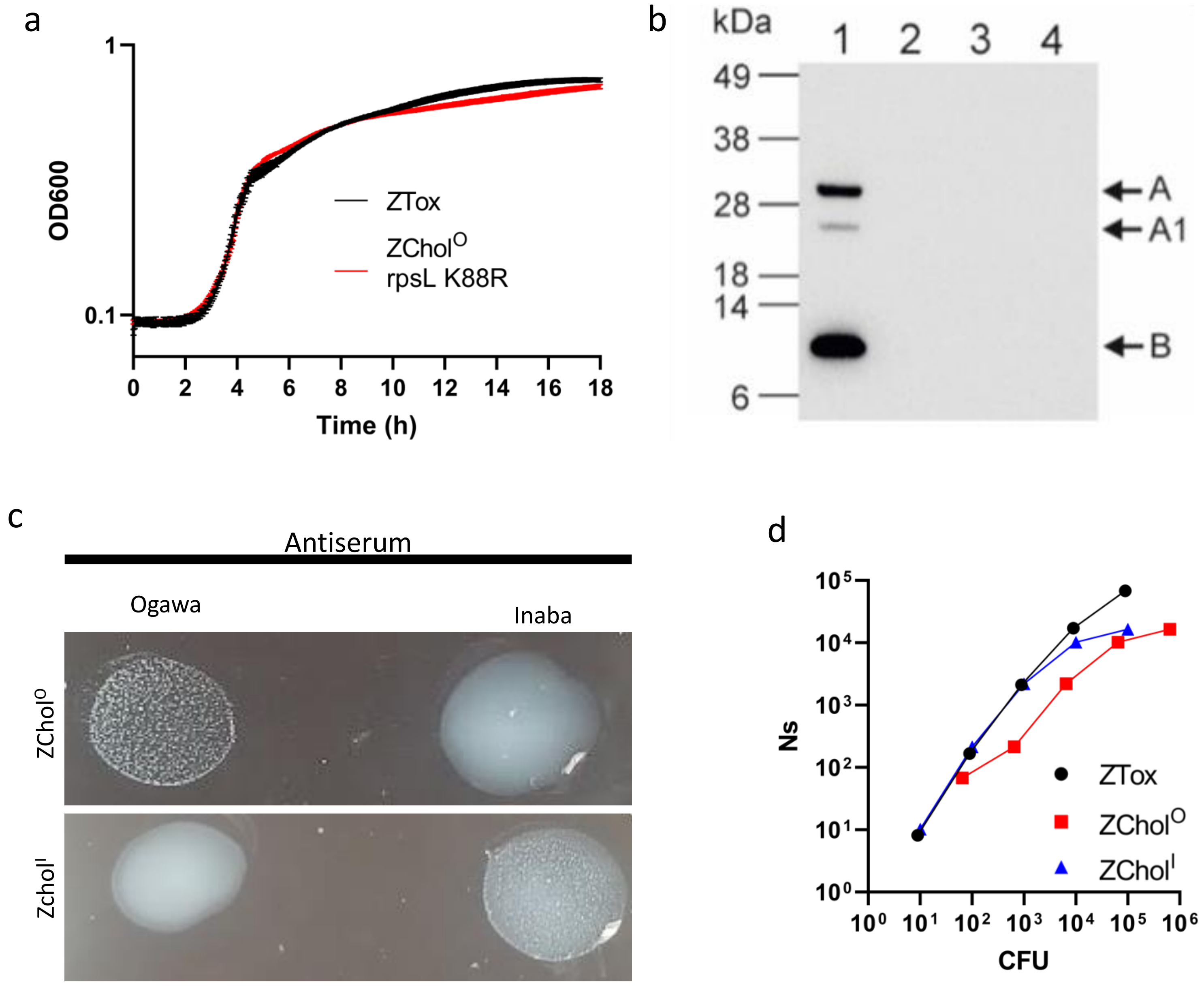
