## Supplemental Table 1 for "Non-toxigenic *Vibrio cholerae* challenge strains for evaluating vaccine efficacy and inferring mechanisms of protection"

| Antibiotic | ZChol$^{O}$ | | ZChol$^{I}$ | |
|---|---|---|---|---|
| | MIC (µg/mL) | Interpretation | MIC (µg/mL) | Interpretation |
| Ampicillin | 4 | S | 4 | S |
| Azithromycin | 0.5 | S | 0.5 | S |
| Ciprofloxacin | 0.5 | S | 0.5 | S |
| Erythromycin | 2 | N | 2 | N |
| Tetracycline | 1 | S | 1 | S |
| Sulfamethoxazole/Trimethoprim | 0.25 | S | 0.25 | S |
| Streptomycin | >200 | R | >200 | R |
